# Repetition effects reveal the sub-sequence representation of actions

**DOI:** 10.1101/2024.08.07.607016

**Authors:** Mahdiyar Shahbazi, J. Andrew Pruszynski, Jörn Diedrichsen

**Affiliations:** Western Centre for Brain and Mind, Western University, London, Ontario, Canada; Department of Statistical and Actuarial Sciences, Western University, London, Ontario, Canada; Department of Computer Science, Western University, London, Ontario, Canada; Department of Physiology and Pharmacology, Western University, London, Ontario, Canada

**Keywords:** Repetition effects, Skill learning, Sequential movements, Memory trace

## Abstract

When a movement sequence is repeated, the second execution is faster than the first. This demonstrates that the brain retains some trace of the just-executed sequence, the earliest form of sequence memory. Currently, it is unclear whether this memory trace is represented at the level of (a) transitions between movements, (b) chunks of multiple movements, or (c) the entire sequence. To answer this question, we instructed human participants to generate sequences of 11 finger presses in a delayed response paradigm. From one trial to the next, segments of variable length (1, 2, 4, 6, 11) could be repeated from the previous trial. We observed that repetition benefits appeared when a segment of 4 consecutive finger presses or longer was repeated from the previous trial. This suggests that the benefit of repetition is not merely the sum of improvements in individual transitions, nor does it require the entire sequence to be repeated. The repetition benefit was small for the first transition of a repeated segment and increased with additional repetitions. This suggests that the memory supporting the repetition effect is mainly activated when a series of past movements matches the memory trace. Planned future movements had less of an effect on the repetition effect. Our results provide insight into the structure of the earliest memory traces for motor sequences.

**Significance Statement:** Many motor skills involve concatenating single movements into complex sequences. Already after a single execution of a sequence, humans retain a memory trace that accelerates the execution of the sequence if it is repeated in the next trial. Our results show a repetition benefit even when only a small portion of movements from the previous trial is repeated, suggesting that full sequence repetition is not necessary. Detailed analyses indicate that the memory trace is activated when the last 2-3 movements match the current execution. Our work, therefore, sheds light on the structure of the earliest sequence memory and will interest researchers studying motor skill learning and the neural basis of movement control.

## Introduction

The best way to improve a motor skill is through repeated practice. Even after a single trial, the human motor system shows some improvements. For example, the execution of a specific sequence causes the following execution of the same sequence to be faster (Ariani et al., 2020), even if there is sufficient time to fully preplan each sequence. The representation remaining in the brain after a single execution is the very first memory trace of a skill and likely forms the seed for a longer-lasting memory representation. Understanding the structure of this initial memory trace, therefore, may offer new insights into how sequence learning occurs.

The control of movement sequences itself has been shown to evoke a hierarchy of representation, ranging from elementary movements to the entire sequence (Ariani et al., 2023; Russo et al., 2020; Shima et al., 2006; Tanji and Shima, 1994; Yokoi and Diedrichsen, 2019). Sequence repetition must facilitate some level of this hierarchy (Berlot et al., 2021), but whether repetition acts at the level of the entire sequence, small sequence components, or individual transitions remains unclear.

To address this question, we designed an experiment asking human participants to generate random sequences of 11 finger movements in a delayed response paradigm. From trial to trial, a variable number of digits (0, 4, 6, 11) could be repeated. The repeated digits could occur consecutively within a segment or break down into single digits or pairs of digits. This allowed us to investigate whether repetition benefits occur by improving the execution of single digits, transitions between digits, some sub-component of the sequence (sub-sequence), or the entire sequence, providing insights into how sequences are represented in the brain. For sub-sequences, we were also able to test whether past and/or future transitions were taken into account to yield a repetition benefit on a repeated transition, thereby providing some insight into the temporal integration window of sequence representations.

## Methods

### Participants

A total of 35 individuals (22 female, mean age = 23±4) took part in the experiment. All participants were right-handed and reported no history of psychiatric or neurological disorders. Participants provided written informed consent for all procedures and data usage before the study started, and all the experimental procedures were approved by the Human Research Ethics Board at Western University. Five participants withdrew from the experiment, and their sessions were terminated before completion. Consequently, their data were excluded from successive analyses (final N = 30, 20 female, age = 24 ± 4).

### Apparatus

Finger presses were produced on a custom-made keyboard with five 10.5 x 2 cm keys. Each key had an indentation to guide fingertip placement. Finger presses were isometric. Forces were measured by transducers (FSG-15N1A; Sensing and Control Honeywell; the dynamic range of 0–25 N; update rate 5 ms) located beneath the fingertip indentation of each key. Five white lines were displayed on a computer screen such that the vertical position of each line was proportional to the force exerted by each finger on the respective key. To register a key press, the applied force had to exceed a 1 N threshold, indicated by a horizontal white line in the middle of the screen (Fig. 1A).

**Figure 1.**
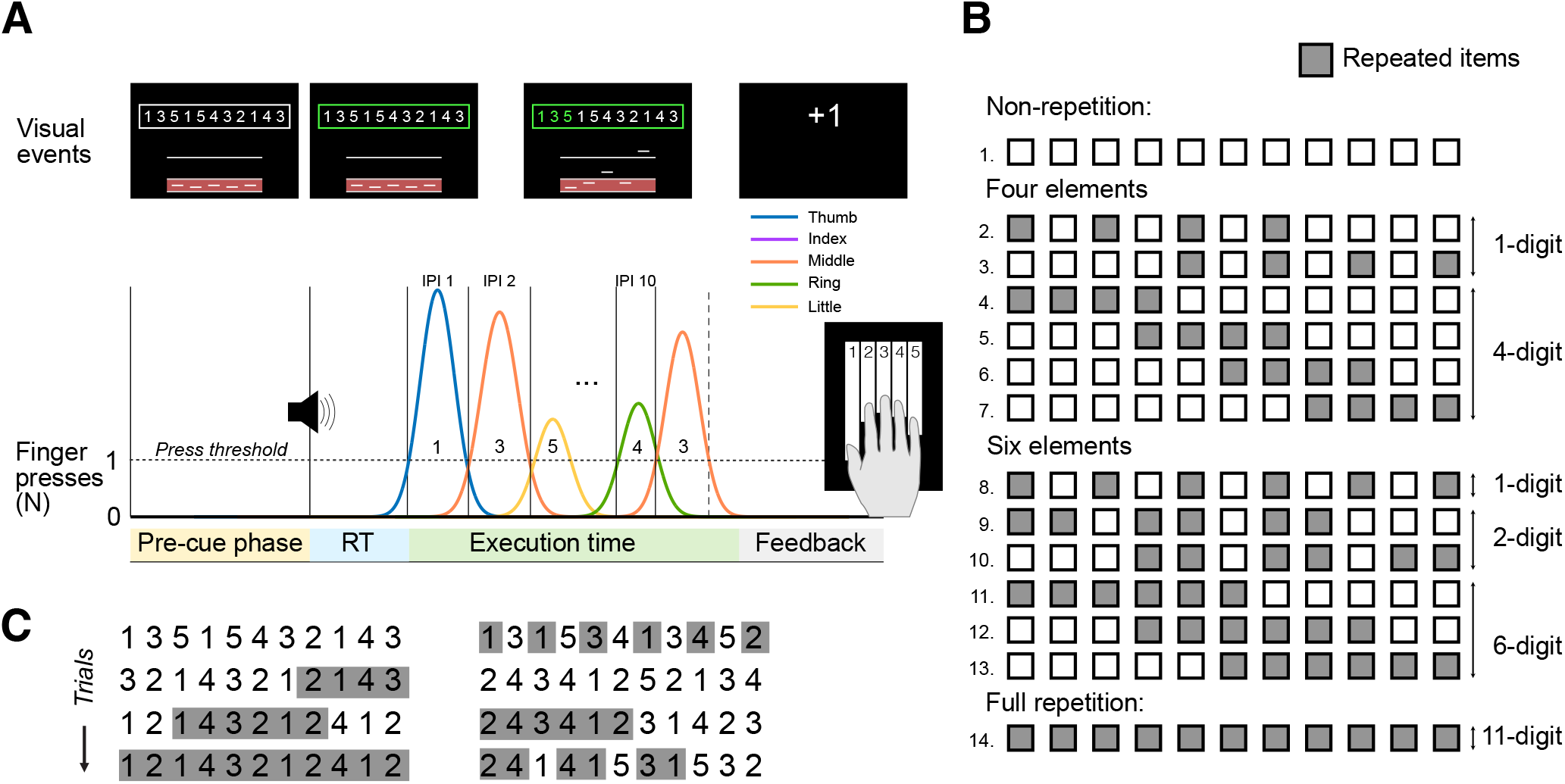
Task design. **(A)** Temporal structure of a trial. In the pre-cue phase (3-4 s), a random sequence of 11 numbers was displayed within a box at the top of the screen. A tone and a color change of the box frame to green then provided a go-signal to execute the sequence as fast as possible. Each correct press caused a digit to turn green. After execution was complete, participants received feedback on their performance for 0.5 s. **(B)** Experimental conditions. Each square represents a single press. A gray square indicates that a digit repeated exactly in that sequential position from the previous trial – a white square indicates a change. Either 0 (Non-repetition), 4 (Four Elements), 6 (Six Elements), or the entire sequence (Full repetition) could repeat. Repeated digits could occur in continuous chunks (4-digit or 6-digit) or divided into pairs (2-digit) or single items (1-digit). **(C)** Example of 8 consecutive trials shown in two columns (trials 1-4 in the left column and trials 5-8 in the right column) with repeated elements highlighted in gray.

### General procedure

We used a discrete sequence production (DSP) task in which participants produced sequences of 11 keypresses with the five fingers of their right hand (Fig. 1A). Each trial was cued by a set of 11 numbers instructing which finger had to be pressed (e.g., 1 = thumb, 2 = index, … 5 = little) in which order. The sequence had to be produced by pressing the fingers corresponding to the numbers, from left to right, as fast as possible. In the pre-cue phase, participants were asked to prepare for the corresponding finger presses. After a random delay of 3-4s, a go-cue marked the beginning of the movement phase. The go-cue was a green frame accompanied by a tone (Fig. 1A), indicating that participants had to perform the planned sequence of finger presses as quickly and accurately as possible.

Performance was evaluated in terms of both execution speed and press accuracy. Speed was defined in terms of total time (TT), which consisted of the reaction time (RT; from the onset of the sequence cue to the first keypress) plus the movement time (ET; from the onset of the first keypress to the release of the last keypress). A single press error invalidated the whole trial, so accuracy was calculated as the percent error rate per block of trials (number of trials with at least one error/number of total trials x 100). In a 500-ms feedback interval, participants were presented with performance points: -1 points for not completing the sequence within 10s; 0 points for ET >5s or for pressing any wrong key; +1 points for correct execution blow 5s; and +3 points for correct execution below the current TT threshold. TT threshold decreased by 2% from one block to the next if both the median TT in the current block was faster than the best median TT recorded hitherto and the mean error rate in the last block was below 20%. If either one of these criteria was not met, the thresholds for the next block remained unchanged. After each block of trials, the median TT, mean error rate, and points earned were displayed to the participants. From block to block, we instructed participants to try to go faster if their error rate was below 15% and to try to be more accurate if their error was above 15%.

### Experimental design

The experiment consisted of two sessions conducted on two consecutive days. In the first session, participants were introduced to the task with 1 training block of 30 trials. Then, participants completed 8 blocks of 60 trials in the first session and 10 blocks of 60 trials in the second session. Each session took ∼120 minutes.

On each trial, participants were presented with a random sequence of 11 numbers. Each number (1-5) was guaranteed to be included at least once but no more than three times in the sequence. Sequences were not allowed to have consecutive runs of three presses (e.g., 1-2-3) or repetitions of a press (e.g., 2-2).

In the next trial, 0, 4, 6, or all elements could repeat. Repeated items could happen successively, as in the “4-digit” or “6-digit” conditions, or in discontinuous pairs or single items, as in “2-digit” or “1-digit” conditions, respectively. Moreover, partial repetitions could occur at the beginning, middle, or end of the sequence, resulting in a total of 12 partial repetition conditions (Fig. 1B,C).

From trial to trial, there was a 20% probability for non-repetition trials and a 20% probability for full repetition trials. This high proportion of non-repetition and full repetition trials allowed us to estimate the baseline performance with high accuracy. Each of the 12 partial repetition conditions occurred with a 5% probability.

### Analysis of reaction and execution times

Within each participant, error trials were removed, and the mean RT and ET were calculated for all 14 conditions across trials and two sessions.

To assess the effect of repetition on RT, we only considered conditions with repeated digits at the beginning of the sequence, such as conditions 4 or 11 (Fig. 1B). A two-tailed paired-sample t-tests was used to compare repetition versus non-repetition conditions.

To assess the effect of repetition on ET, we averaged conditions with the same number of repeated fingers and equal length of the repeated segment, differing only in the location of repetition (e.g., conditions 11, 12, 13 for 6-digits). Statistical analysis for assessing the effect of ET, independent of location, included two-tailed paired-sample t-tests comparing repetition versus non-repetition conditions. We also used a within-subject repeated measures ANOVA to assess the effect of location on repetition.

### Analysis of inter-press intervals

For a finer-grained analysis of the repetition effect, we also analyzed the specific inter-press intervals (IPIs) following repetition. An IPI was defined as the time between subsequent finger presses, i.e., the time between the two subsequent fingers crossing the 1N threshold (Fig 1A). After removing error trials, we averaged the IPI for each transition (10 transitions) and condition (14 conditions) within each participant (over trials of two sessions), resulting in 140 values. For each position in the sequence (1-10), we then calculated the difference between each condition and the corresponding non-repetition IPI, resulting in 130 values per participant. This allowed us to measure the repetition benefit independent of the baseline time required to complete a transition. To summarize the data, we grouped the transitions as follows: “*pre*” if a non-repeated transition occurred before a repeated transition, “*rep*” if the transition was repeated, “*post*” if a non-repeated transition occurred after a repeated transition, and “*nrep*” otherwise. Furthermore, we grouped “*rep*” transitions into “*first*” if it was the initial transition within a repeated segment, “*last*” if it was the last one, and otherwise as “*middle*”. This grouping was conducted separately for 2-digit, 4-digit, and 6-digit conditions.

We used two-tailed paired-sample t-tests to assess the repetition changes in IPI groups. Additionally, we used within-subject repeated measures ANOVAs to assess if repetition changes vary between the first, middle, and last IPIs.

### Modelling repetition changes

To characterize the repetition effects across all conditions, we tested a series of general linear models of the observed repetition in each of the 130 possible IPIs. The simplest model, termed the “*current transition model*” (or “curr” for short), included a single feature indicating whether the current transition was repeated or not. We compared the predictive power of this model with three more complex models. The “*current + past transition model*” (“curr+past” for short) contained three features: whether the current transition was repeated, whether the previous transition was repeated, and whether both were repeated (the interaction term). The “*current + next transition model*” (“curr+next” for short) included features for the current and the upcoming transitions, as well as their interaction. Finally, the “*current + past + next transition model*” (“curr+past+next” for short) included features for the current, previous, and upcoming transitions, as well as all possible two-way interactions among these. All models also included an intercept term.

To compare the models, we estimated the predictive R^2^ using a 10-fold cross-validation scheme. We first concatenated the data of all subjects into a single data vector (130 samples per subject × 30 subjects) and then divided it into 10 random folds. For each model, we estimated the linear regression parameters using the 9 folds and calculated the R^2^ on the 10^th^ fold. We then computed the mean R^2^ over all 10 iterations. We used the same folds for all models.

We then compared each model to the “current transition model” by taking the difference in their predictive R^2^ term. To find a 95% confidence interval for this difference, we repeated the same process 15,000 times by bootstrapping over participants. If the lower bound of the interval was above zero, then that model was considered to have a greater predictive power than the current transition model.

## Results

### Participants benefit from repeating a sub-sequence in long movement sequences

We first asked whether the sequence repetition benefit can arise when only part of the sequence is repeated, and what length of repetition is necessary to observe this benefit. To do so, we varied the number of repeated digits, repeated transitions, and the length of repeated subsequences embedded in otherwise random sequences (Fig. 1B, C).

When the entire sequence was repeated, the execution time was 125 ± 16 ms faster than in the non-repetition condition (Fig. 2A; *t*_*29*_ *= 7.731, p = 1.5e-8*). For partial repetitions, we only found an overall effect on ET when participants repeated 6 consecutive digits from the previous trial (Fig. 1B, conditions 11, 12, 13 averaged; 51 ± 14 ms, *t*_*29*_ *= 3.670, p = 9.7e-4*). This partial repetition benefit did not depend on the placement of repeated items – the repetition benefit did not vary whether it occurred at the beginning, middle, or end of the sequence (repeated-measures ANOVA: *F*_*2*,*58*_ *= 0.492, p = 0.6139*). In contrast, the repetition of 4 consecutive digits (conditions 4, 5, 6, 7 averaged; *t*_*29*_ *= 0.628, p = 0.5347*), the repetition of 6 digits arranged in three 2-digit pairs (conditions 9, 10 averaged; *t*_*29*_ *= 1.275, p = 0.2125*), the repetition of 6 isolated digits (condition 8; *t*_*29*_ *= 0.512, p = 0.6126*), and the repetition of 4 isolated digits (conditions 2, 3 averaged; *t*_*29*_ *= -0.649, p = 0.5213*) did not lead to a significant overall effect on ET. Together, these findings suggest that the repetition benefit does not require the entire sequence to be repeated but can occur with a sub-sequence of at least length 6. The repeated sub-sequence can be reused flexibly, independent of its location within the sequence.

In agreement with our previous report (Ariani et al., 2020), we also found a small repetition effect on RT in the full repetition condition (10 ± 3 ms, *t*_*29*_ *= 3.146, p = 0.003*). No statistically significant effect was observed in the other conditions, even if we restricted the analyses to conditions in which the repeated digits were at the beginning of the sequence (Fig. 2B; *t*_*29*_*<1.882, p>0.0699*).

**Figure 2.**
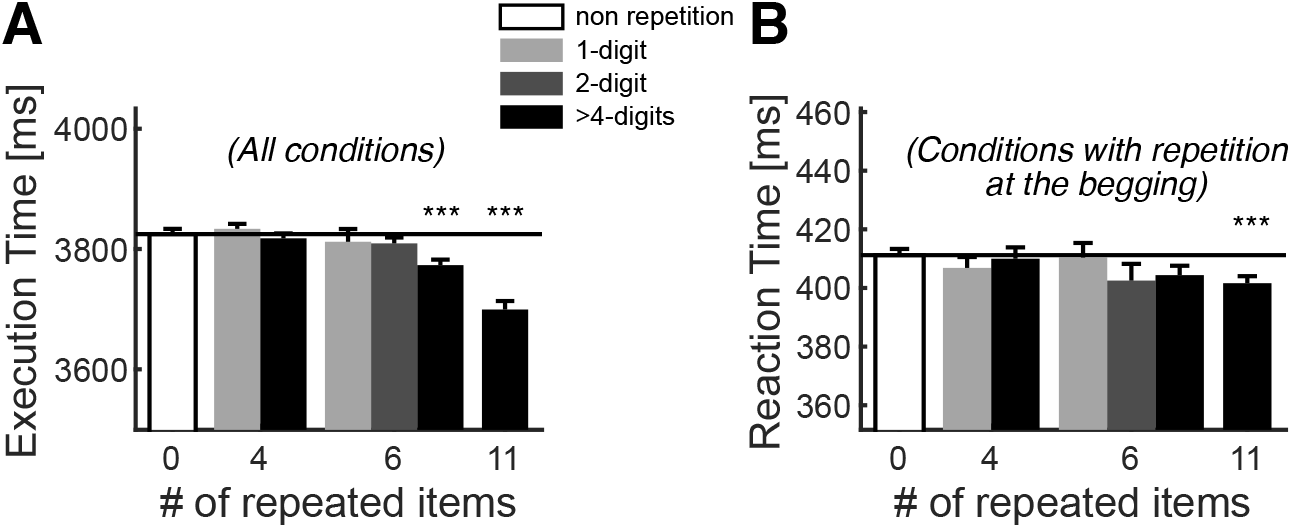
Sequential nature of repetition improvement. **(A)** Group-averaged execution times for all conditions, categorized by the number of repeated digits, are presented. The contrasts indicate the length of the repeated sub-sequence (black: 4, 6, and 11-digit sequence; dark gray: 2-digit transitions; light gray: isolated repeated digits). The solid black line represents the execution time for the condition with no repetition. Error bars represent the SEM across participants. ***p<0.001, using two-tailed paired-sample t-tests of each condition against the non-repeated condition. **(B)** Group-averaged reaction times are shown in the same format as in (A) but for conditions with repetitions occurring at the beginning of the sequence.

### The activation of a memory trace depends on both current and previous transitions

The overall execution time only improved following the repetition of 6 or 11 consecutive items. The lack of a significant effect when 1, 2, or 4 consecutive finger presses were repeated, however, could reflect that the repetition introduced both speed-ups and slow-downs at different parts of the sequence in such a way that the overall execution time did not statistically improve. To investigate this more closely, we performed a detailed analysis of the inter-press intervals (IPIs).

We first analyzed the IPIs for full and non-repetition conditions across the entire sequence (Fig. 3A). In both conditions, the middle transitions were slower than the initial and final ones. This pattern is ubiquitous in DSP tasks and can be explained by the fact that the first few items can be pre-planned, enabling fast execution. In the middle of the sequence, participants have to plan the new actions on the fly (online planning), which slows their performance (Ariani et al., 2021). Finally, in the end, more resources are available to plan the last items as no more future items need to be taken into consideration, which again speeds up performance.

**Figure 3.**
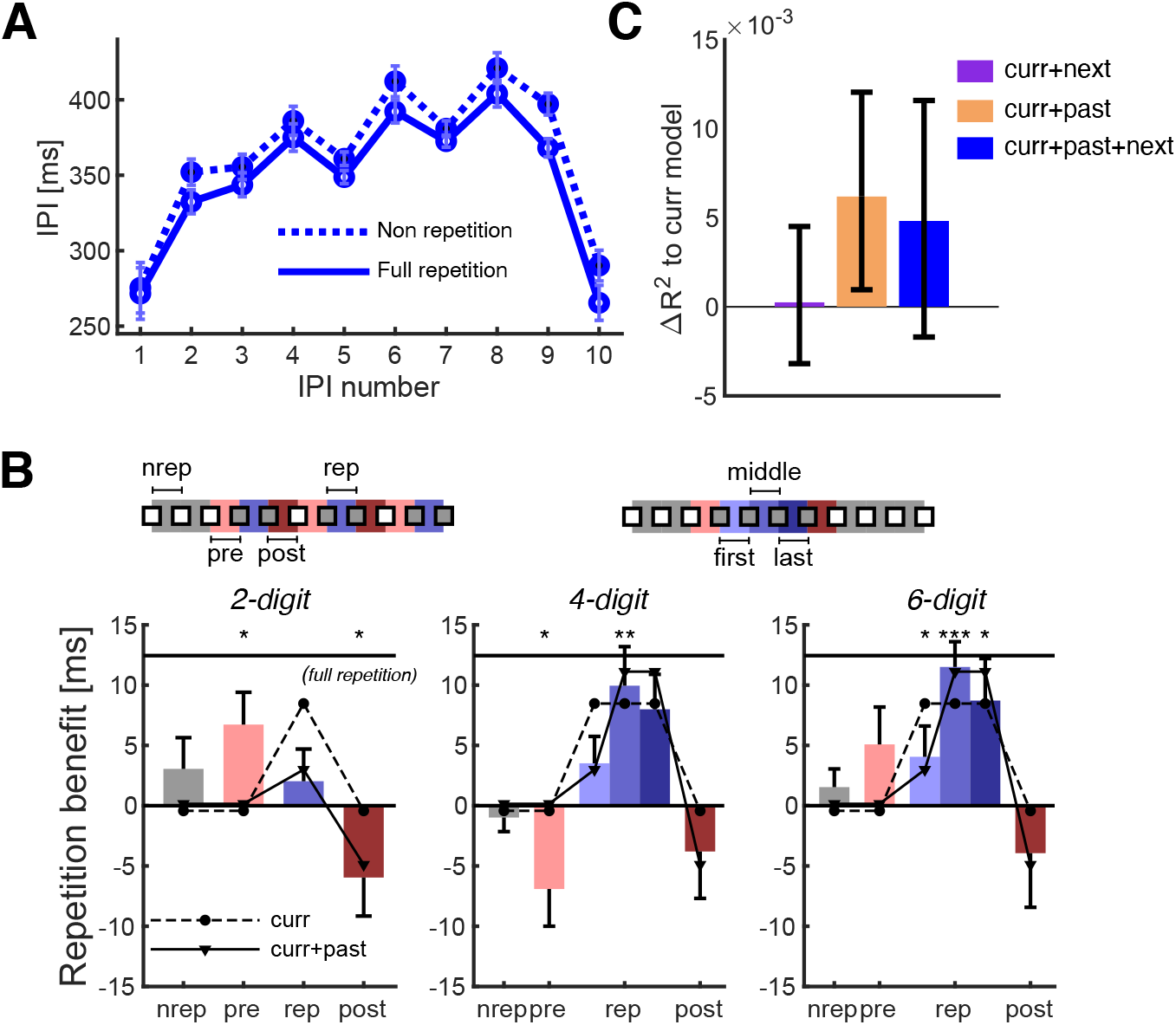
Repetition benefit is not solely an improvement in movement transitions. **(A)** IPIs for full repetition (solid line) and non-repetition (dotted line) conditions as a function of the position in the sequence. **(B)** Average repetition benefit on different transitions relative to the IPI at the same sequential position in the non-repetition condition. The solid line represents the magnitude of the benefit in the full repetition condition. IPIs are grouped as “pre” (light red) if a non-repeated IPI occurred before a repeated IPI, as “post” (dark red) if it occurred after, as “nrep” (gray) if it was not associated with repeated digits, and as “rep” (blue) if it was repeated. The repeated transitions were further grouped as “first,” “middle,” or “last,” depending on their position in the repeated subsequence. The dashed lines indicate the fit of the curr model, and the solid lines indicate the fit of the curr+past model. Error bars indicate the SEM across participants. *p<0.05, **p<0.01, ***p<0.001, two-tailed paired-samples t-test. **(C)** Difference of R^2^ of curr+next (purple), curr+past (orange), and curr+past+next (blue) from the curr model.

For each of the transitions, we observed a small repetition benefit. Consistent with a previous report (Ariani et al., 2020), the repetition benefit was smaller in the first transition and larger later in the sequence, suggesting that the repetition benefit arises from an acceleration of online planning. To analyze this repetition benefit across conditions, we used the non-repetition IPI for each sequential position as a baseline and subtracted it from the IPI data for each participant (see Methods).

For 1-digit conditions, none of the IPIs were faster than the corresponding IPI in the non-repetition conditions (*t*_*29*_*<1.741, p > 0.0923*), suggesting that having an isolated digit at the same position as in the last sequence is not sufficient for a repetition effect. For repeated transitions (2-digit conditions), we also did not find a repetition benefit (Fig. 3C, left; *t*_*29*_ *= 0.725, p = 0.4744*).

For 4-digit conditions, however, a 7 ± 2 ms repetition benefit was observed on repeated transitions (Fig. 3C, averaged over the first, middle, and last repeated transitions; *t*_*29*_ *= 4.182, p = 2.4e-4*). As expected, this was also true for 6-digit conditions (8 ± 2 ms, *t*_*29*_ *= 4.959, p = 2.8e-5*). The repetition benefit in the middle transition was nearly the size of the full repetition condition. Thus, a repetition benefit can occur when the repeated segment is at least four digits long.

Interestingly, in the 4 and 6-digit conditions, the speed-up in the first repeated transition was by 7±3 ms smaller than in the middle transitions (*t*_*29*_ *= 2.745, p = 0.010*). In other words, the repetition effect was stronger when both the current and previous transition matches the memory trace, suggesting that the memory requires some time to be activated.

In contrast, the repetition effect did not seem to be sensitive to the next transition. For the last repeated transition in repeated sub-sequence, the repetition benefit was as large as in the middle of the subsequence (*t*_*29*_ *= 0.795, p = 0.4331*). Therefore, the influence of the memory trace does not seem to diminish, even when a future planned transition does not agree with the memory.

Finally, in the first non-repeated transition after a repeated subsequence we found a significant -5±2 ms slow-down (2, 4, and 6-digit conditions averaged; *t*_*29*_ *= -2.238, p = 0.0330*). One reason for this consistent slow-down could be a mismatch between an activated memory trace and the incoming sensory information, which requires some time to be resolved.

To systematically test for the contribution of the current, previous, and upcoming transitions to the repetition changes, we built four models incorporating different combinations of these factors (see Methods). The baseline (‘curr’) model only considered whether the current transition was repeated without any look-back or look-ahead window. Naturally, this model predicted equal benefits for all repeated IPIs (Fig. 3C, dashed line). The predictive R^2^ value of this model was 0.0127.

We then asked whether adding the information about the previous transition (curr+past model) resulted in a better prediction of repetition benefit. This model predicted smaller benefits for the first transition as well as post-repetition slow-down (Fig. 3C, solid lines). We found that the predictive R^2^ of this model was significantly larger than the baseline model by CI_95_= [0.001, 0.012] (Fig. 3B).

The curr+past model did not predict the slight slow-down in the last transition compared to the middle one (Fig. 3C). Naturally, the model containing the information of the upcoming transition (curr+past+next model) was able to do that. However, adding information about the next transition to the curr and curr+past models did not result in a significant increase in predictive power. This suggests that the importance of the previous transition is more pronounced than that of the future one in activating the memory trace following sequence repetition.

## Discussion

Our study investigated the structure of the first memory trace of a motor sequence, revealing key insights into how the brain processes and retains sequential information. Previous work has shown that, even after a single execution of a new sequence, the next execution of the same sequence is faster(Ariani et al., 2020; Johnson et al., 2023). However, it remained unclear whether this improvement is due to the facilitation of individual movements (Mawase et al., 2018), the transitions between them, chunks (or sub-sequences) of three or more movements, or the entire sequence.

In our study, we show that the repetition benefit does not only occur when the entire sequence is repeated. Even when only a sub-sequence of four consecutive movements was repeated from the preceding sequence, the execution of those finger presses was faster. This phenomenon has also been observed in multi-day sequence learning studies, in which a learned chunk (or a sub-sequence) of 2-3 movements could be executed faster, even if it was embedded in a random long sequence (Sakai et al., 2003; Verwey, 2001; Wong et al., 2015; Yokoi and Diedrichsen, 2019). A more recent study looking at the generalization of learning after a single trial (Johnson et al., 2023) failed to find a benefit of a repeated subsequence of 3 presses. However, with careful counterbalancing of sequences and detailed analysis of different inter-press intervals, our findings indicate that such benefits are clearly present.

Thus, single-trial repetition and long-term motor skill learning appear similar in terms of behavioral generalization. This suggests that the initial sequential memory trace may have similar structure to that of long-term memory. Previous studies (Ariani et al., 2020; Berlot et al., 2021) have also shown that the repetition benefits decrease as long-term learning progresses. This indicates that sequence repetition acts on the same representation that improves during multi-day learning.

The neural mechanisms underlying this phenomenon likely involve the premotor and superior parietal areas of the brain. Notably, these regions have been shown to exhibit repetition suppression— reduced fMRI activity upon repeated sequence execution—even when the speed of execution is controlled (Berlot et al., 2021). This suppression occurs in the same areas that are involved in long-term sequence learning, suggesting that the transient memory trace supporting repetition suppression shares a neural substrate with the stable memory trace that supports long-term skill retention (Wiestler and Diedrichsen, 2013; Yokoi and Diedrichsen, 2019). This convergence of short-term and long-term memory processes highlights the role of these brain regions in both immediate and prolonged motor learning.

Our findings suggest that the nervous system breaks long sequences into manageable parts rather than relying on a single, unified representation. This insight is supported by our observation that increasing the length of the repeated segment beyond four movements does not provide any additional repetition benefit (Fig. 3B). However, it is still not clear whether the nervous system subdivides sequences discretely into fixed sub-sequences (i.e., chunks) or if it controls sequences using a finite temporal window, spanning both past and future targets. Dissociating the two possibilities is challenging because we do not know for certain *how* participants will break up a given sequence. First, the boundaries of chunks, indicated by slowdowns during sequence execution, differ among individuals and can change within the same participants during sequence learning (Popp et al., 2020). Second, slowdowns during sequence execution can also occur due to the biomechanical characteristics of the movements. Not knowing for certain where these chunk boundaries fall makes it harder to compare generalization within chunk with generalization across chunk boundaries.

Our findings provide insight into the neural interactions between internal and external sequence representations (Ariani et al., 2015; Honey et al., 2018; Mizes et al., 2023). In our task, participants needed to read the numbers from the screen and convert these external cues into motor commands to execute the sequence. For the repetition benefit to occur, the externally cued sub-sequence must be able to activate the lingering internal representation if it is matching the current demands.

One insight into the interaction between external stimuli and the memory trace comes from the finding that the repetition benefit was smaller at the beginning of the repeated segment and grew as more elements were repeated (Fig. 3B). This result suggests that, although the brain maintains an internal memory representation of all repeated finger presses, this memory is not fully activated until the memory and external cue match for a few finger presses. That is, memory activation appears to depend both on ongoing and previous finger presses.

Another insight comes from the observation that participants slow down, once the repeated sub-sequence ends and an unfamiliar part of the sequence begins (Fig. 3B). Participants execute this press even more slowly than in a non-repeated sequence, suggesting that the slowdown results from interference between the new stimuli and the activated memory representation. This conflict likely creates competition that requires extra time to resolve.

Previous work has shown that the motor system plans a number of upcoming movements while controlling the current movement (Ariani et al., 2021; Kashefi et al., 2024; Shahbazi et al., 2024). Given this, it might be expected that participants slow down once they detect a mismatch between a future stimulus and a future planned movement. This was not the case as the last IPI of a repeated sub-sequence was not significantly slower than the one in the middle of the sub-sequence (Fig 3B). Therefore, the influence of the lingering memory trace appears to be modulated only by the match with past (executed) movements, but not on the match with future (planned) movements. This observation provides important constraints on the neural mechanisms that control the interactions between externally cued sequence execution and the earliest forms of sequence memory.

## Acknowledgments

This work was supported by a CIHR Project Grant to JD and JAP (PJT-175010) and the Canada First Research Excellence Fund (BrainsCAN) to Western University. JAP received a salary award from the Canada Research Chairs program.

